# Relief of the Dma1-mediated checkpoint requires Dma1 autoubiquitination and dynamic localization

**DOI:** 10.1101/322214

**Authors:** Christine M. Jones, Jun-Song Chen, Alyssa E. Johnson, Zachary C. Elmore, Sierra N. Cullati, Janel R. Beckley, Kathleen L. Gould

## Abstract

Chromosome segregation and cell division are coupled to prevent aneuploidy and cell death. In the fission yeast *Schizosaccharomyces pombe*, the septation initiation network (SIN) promotes cytokinesis, but upon mitotic checkpoint activation, the SIN is actively inhibited to prevent cytokinesis from occurring before chromosomes have safely segregated. SIN inhibition during the mitotic checkpoint is mediated by the E3 ubiquitin ligase Dma1. Dma1 binds to the CK1-phosphorylated SIN scaffold protein, Sid4, at the SPB, and ubiquitinates it. Sid4 ubiquitination antagonizes the SPB localization of the Polo-like kinase Plo1, the major SIN activator, so that SIN signaling is delayed. How this checkpoint is silenced once spindle defects are resolved has not been clear. Here we establish that Dma1 transiently leaves SPBs during anaphase B due to extensive auto-ubiquitination. The SIN is required for Dma1 to return to SPBs later in anaphase. Blocking Dma1 removal from SPBs by permanently tethering it to Sid4 prevents SIN activation and cytokinesis. Therefore, controlling Dma1’s SPB dynamics in anaphase is an essential step in S. *pombe* cell division and the silencing of the Dma1-dependent mitotic checkpoint.

## Introduction

Accurate cell division, yielding two genetically identical daughter cells, requires coordination between mitosis and cytokinesis. In the fission yeast *Schizosaccharomyces pombe*, the septation initiation network (SIN), a protein kinase cascade, coordinates these two events by initiating cytokinesis after chromosome segregation (for review see (Krapp and Simanis, 2008; Johnson et al., 2012b; Simanis, 2015)). When there is a mitotic error, a checkpoint mechanism inhibits SIN signaling to prevent cytokinesis from occurring before chromosomes have safely segregated (Marks et al., 1992; Murone and Simanis, 1996; Guertin et al., 2002; Johnson and Gould, 2011). This checkpoint operates in parallel to the spindle assembly checkpoint (Murone and Simanis, 1996; Johnson et al., 2013; Musacchio, 2015). Checkpoints with similar purpose of maintaining coordination between chromosome segregation and cytokinesis also exist in *Saccharomyces cerevisiae* and mammals (Mendoza and Barral, 2008; Caydasi and Pereira, 2012; Nahse et al., 2017).

SIN inhibition during a mitotic error depends on the dimeric E3 ubiquitin ligase Dma1, a member of the FHA and RING finger family (Brooks et al., 2008). Dma1 was identified as a high copy suppressor of a hyperactive SIN mutant and it is required to prevent septation during a prometaphase arrest elicited by the ß-tubulin mutant *nda3-km311* (Hiraoka et al., 1984; Murone and Simanis, 1996). Overproduction of Dma1 completely blocks SIN activity and results in cell death (Guertin et al., 2002), indicating its levels and activity must be properly regulated. Dma1 co-localizes with SIN components both at the cell division site and at the mitotic spindle pole body (SPB) (Guertin et al., 2002), however it is Dma1’s SPB localization and its ubiquitination of the SIN scaffold protein, Sid4 (Chang and Gould, 2000; Morrell et al., 2004), that is required for SIN inhibition during a mitotic checkpoint (Johnson and Gould, 2011). Sid4 ubiquitination antagonizes the SPB localization of the Polo-like kinase Plo1 (Guertin et al., 2002; Johnson and Gould, 2011), the major SIN activator (Ohkura et al., 1995; Mulvihill et al., 1999; Tanaka et al., 2001) so that SIN signaling is blocked and cytokinesis is delayed.

Upon resolution of the mitotic spindle defect, the Dma1-dependent checkpoint signal must be extinguished in order to resume SIN activation and cell division. Here we report that Dma1 exhibits striking fluctuations in its localization to SPBs and the division site during the cell cycle. We found that both SIN activity and Dma1 auto-ubiquitination modulate Dma1 SPB localization dynamics and therefore its ability to inhibit the SIN. By permanently tethering Dma1 to SPBs and preventing these fluctuations, the SIN fails to become active and cytokinesis is blocked. Therefore, Dma1’s dynamic SPB localization is a critical feature of S. *pombe* cytokinesis and silencing of the mitotic checkpoint.

## Results

### Dma1’s E3 ligase activity impacts its abundance and localization

Consistent with previous literature (Guertin et al., 2002; Johnson and Gould, 2011), we observed that Dma1 tagged at its endogenous locus with mNeonGreen (mNG) (Shaner et al., 2013; Willet et al., 2015) (Figure 1A) concentrates at SPBs during mitosis, where it is required for SIN inhibition (Guertin et al., 2002). We also observed localization at cell tips during interphase and at the division site during mitosis and cytokinesis. Given that wildtype Dma1 did not appear to localize to SPBs during the majority of interphase (Figure 1A), we sought to identify the mechanism(s) that influences Dma1 SPB targeting and SIN inhibition.

**Figure 1.**
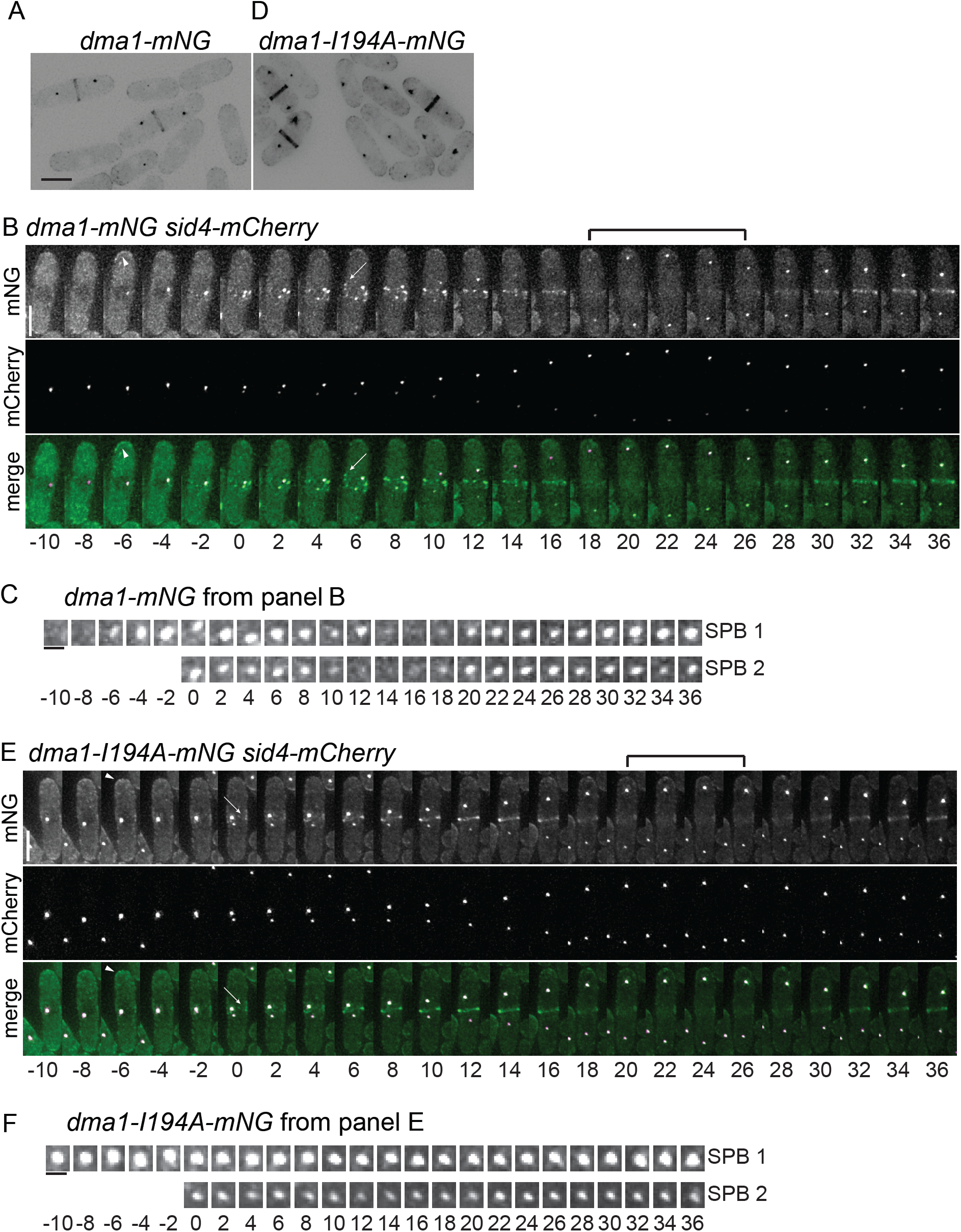
Dynamics of Dma1 localization through the cell cycle. (A and D) Live cell images of *dma1-mNG* (A) and *dma1-I194A-mNG* (D). Scale bar, 5 μm. (B and E) Images from representative movies of *dma1-mNG sid4-mCherry* (B) and *dma1-I194A-mNG sid4-mCherry* (E). Arrows indicate Dma1 localization in cytokinetic node-like structures. Arrowheads indicate Dma1 localization to cell tips. Brackets indicate times of reduced Dma1 detection at the division site. Time in minutes denoted below images; 0 indicates initial frame of SPB separation. Scale bars, 5 μm. (C and F) Enlarged SPB region(s) from movies in B and E. Scale bars, 1 μm.

Because Dma1 localization changes over the course of the cell cycle, we undertook time-lapse imaging experiments to clarify the timing of Dma1 localization to the SPB and cell division site. Dma1-mNG became enriched at SPBs prior to SPB separation (Figures 1B and S1A). Unexpectedly, at the onset of mitosis, Dma1-mNG appeared in node-like structures, a pattern previously undetected for Dma1 (Guertin et al., 2002) and reminiscent of cytokinetic precursor nodes (Rincon and Paoletti, 2012), before forming a ring at the division site (Figure 1B). Then, Dma1-mNG appeared to transiently leave SPBs during anaphase B, returning to them before telophase and then leaving again after cell division (Figures 1B, 1C and S1B). In 24 out of 28 (86%) mitotic SPBs, Dma1-mNG signal transiently dimmed or became undetectable whereas the SPB markers Sid4-mCherry or Sad1-mCherry did not. Dma1 SPB dimming occurred within 2 minutes of anaphase B onset in 90% of cells exhibiting this phenomenon, and it returned to SPBs in all cases before the end of anaphase B, marked by maximal SPB separation, and then left again at cell division. In a few cases, the Dma1 signal was observed to “flicker” on and off a SPB during anaphase B and to shift in intensity between the two SPBs.

Similarly, Dma1-mNG was observed to leave and then return to the division site although with delayed kinetics (mins 18–26) compared to what was observed at SPBs (mins 14–16) (Figures 1B and C, S1A and B). The role of Dma1 at the division site is not yet understood because no binding partners or substrates at the medial cortex have been identified.

Interestingly, in contrast to wildtype Dma1, we found that the Dma1-I194A mutant, which lacks ubiquitin ligase activity due to a mutation in the RING-finger domain (Johnson and Gould, 2011), was detectable at SPBs throughout the cell cycle when it was tagged with mNG (Figure 1D) or GFP (data not shown) at its C-terminus, or either fluorophore at its N-terminus (data not shown). Dma1-I194A-mNG signal persisted at SPBs throughout the cell cycle, only detectably dimming at 14% of mitotic SPBs, although its dynamic localization at the cell division site appeared like wildtype (Figures 1E and S1C). Also, in 20 of 20 cells Dma1-I194A-mNG localized more intensely at one of the two SPBs for most of mitosis (Figure 1E and F). Thus, Dma1 localization is more dynamic than previously appreciated, and Dma1 catalytic activity affects its SPB dynamics. Because Dma1 activity does not impact its cell division site localization dynamics, a different mechanism of targeting to this site must be in place.

### Sid4 ubiquitination does not impact Dma1 accumulation at SPBs

To determine why Dma1-I194A was less dynamic at SPBs during mitosis than Dma1, we examined whether the ubiquitination status of either of Dma1’s two known substrates, Sid4 and Dma1 itself (Johnson and Gould, 2011), modulated its localization. To investigate if the absence of Sid4 ubiquitination promoted Dma1 localization to SPBs, we first needed to develop a strain in which Sid4 ubiquitination was abrogated. The canonical approach is to replace target lysines with arginines; however, previous attempts to construct such a Sid4 variant were unsuccessful (Johnson and Gould, 2011; Johnson et al., 2013). We therefore turned to a method described previously to eliminate ubiquitination of a protein of interest—fusion to a deubiquitinating enzyme (DUB) catalytic domain (Stringer and Piper, 2011).

To identify the appropriate DUB for fusion to Sid4, we screened eighteen of the twenty *S*. *pombe* DUBs for their ability to rescue Dma1 overexpression-induced cytokinesis failure and cell death (Murone and Simanis, 1996; Guertin et al., 2002). Four of the eighteen DUBs (Ubp1, Ubp2, Ubp7, and Ubp14) suppressed Dma1-induced cell death, presumably by reversing Sid4 ubiquitination. Of these, only Ubp7 has diffuse cytoplasmic localization and functions independently of other subunits *in vivo* (Kouranti et al., 2010), making it well-suited for our purpose. The ubiquitin specific protease (USP) domain of Ubp7 was fused to the C-terminus of Sid4 (Sid4-DUB) and produced under control of the native *sid4+* promoter as the sole version of Sid4 in the cell. The fusion did not affect cell viability, but the Sid4-DUB fusion was still ubiquitinated (Figure S2A), indicating that the DUB was not able to access Sid4 ubiquitination sites.

We next tested whether adding the Ubp7 USP domain to the C-terminus of the Sid4 binding partner Ppc89 (Rosenberg et al., 2006) eliminated Sid4 ubiquitination. The Ppc89-Ubp7 USP fusion (hereafter called Ppc89-DUB) abolished Sid4 ubiquitination comparable to deletion of *dma1* (Figure S2B). The *ppc89-DUB* strain grew similarly to wildtype at a variety of temperatures (Figure S2C), and as would be expected when Sid4 cannot accumulate ubiquitin modifications, the *ppc89-DUB* strain resisted Dma1 overexpression-induced cell death (Figure S2D).

To determine if lack of Sid4 ubiquitination affected Dma1-mNG localization, we measured and compared Dma1-mNG SPB intensity relative to Sad1-mCherry in *wildtype* and *ppc89-DUB* strains and found no difference (Figure S2E and F). Moreover, the dynamic localization of Dma1-mNG to the SPB and division site was unchanged in the *ppc89-DUB* strain, although mitotic progression took longer in this strain; of 22 SPBs examined in 11 cells, Dma1-mNG was transiently undetectable on 17 and diminished on 5 others during anaphase (Figure S2G and H). These data demonstrate that an absence of Sid4 ubiquitination does not account for the differences observed in catalytically inactive Dma1 dynamic localization at SPBs relative to wildtype Dma1.

### Dma1 exhibits promiscuous auto-ubiquitination *in vivo* and *in vitro*

In addition to displaying distinct dynamics, by comparing Dma1-mNG and Dma1-I194A-mNG intensities normalized to the SPB marker Sad1-mCherry (Hagan and Yanagida, 1995), we found that Dma1-I194A-mNG was more abundant (3.2-fold) at SPBs in both mitotic and septated cells compared to wildtype Dma1 (Figure 2A). Although we did not quantitate Dma1-I194A abundance at the division site or cell tips, it was visibly more intense than wildtype Dma1 at these sites as well (Figure 1D).

**Figure 2.**
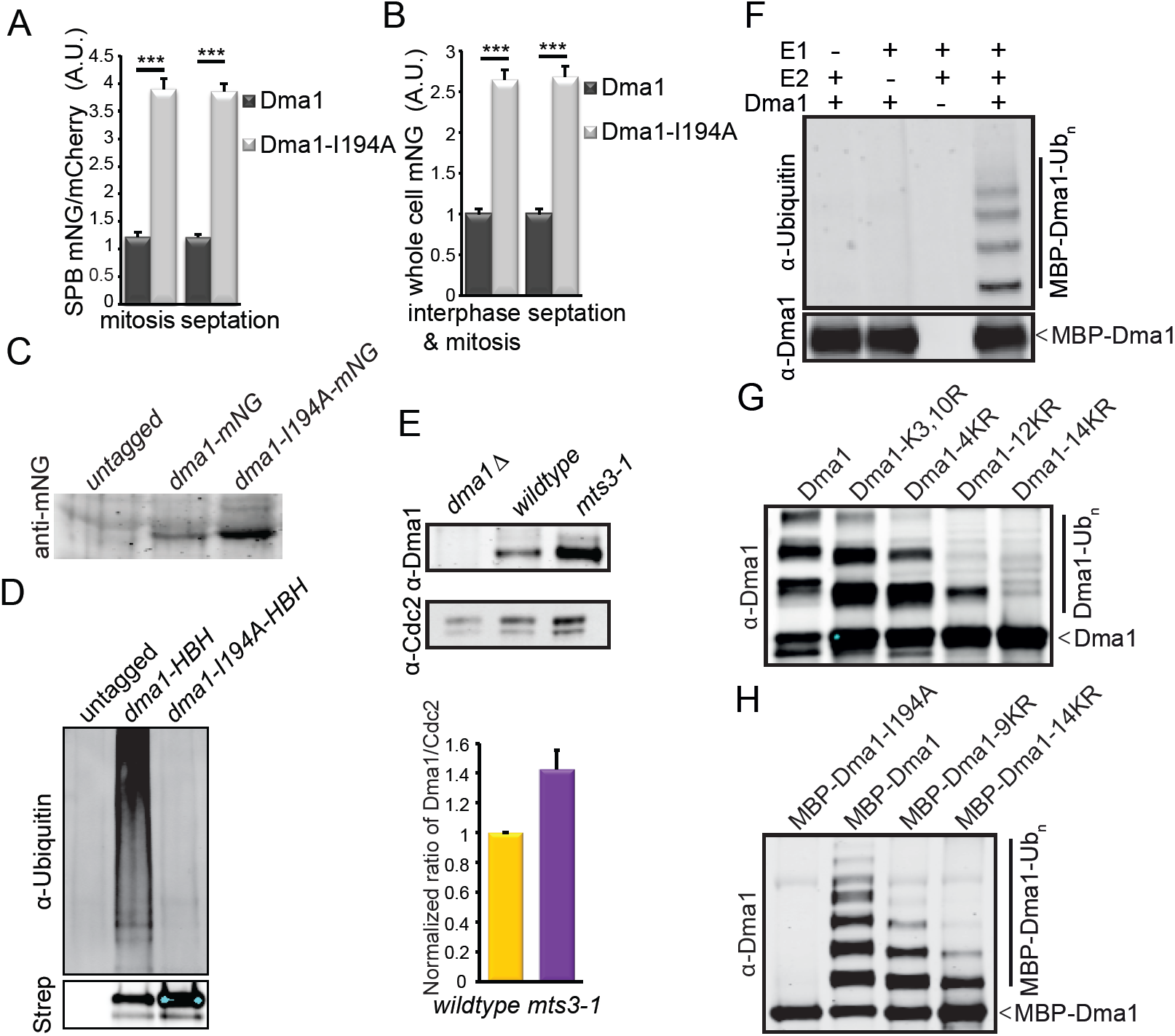
Dma1 auto-ubiquitination influences its abundance and localization dynamics. (A) Quantification of Dma1-mNG and Dma1-I194A-mNG intensities at SPBs, relative to Sad1-mCherry in mitotic or septated cells. n ≥ 42 cells for each measurement; error bars represent standard error determined by two-tailed Student’s t-test, ***p = 4.9×10^−43^ (mitosis) and 1.3×10^−11^ (septation). A.U. = arbitrary units. (B) Quantification of Dma1-mNG and Dma1-I194A-mNG whole cell fluorescence intensities in non-septated interphase and mitotic cells, or septated cells. n ≥ 20 cells for each measurement; error bars represent standard error determined by two-tailed Student’s t-test, ***p =1.3×10^−7^ (interphase and mitosis) and 4.9×10^−9^ (septation). A.U. = arbitrary units. (C) Abundance of Dma1-I194A-mNG relative to wildtype Dma1-mNG was determined by immunoblotting. One representative blot of 3 independent repetitions is shown. (D) Dma1-HBH, Dma1-I194A-HBH, or non-specifically purified proteins were isolated from *mts3–1* cells that had been shifted to 36°C for 3 hr. Dma1 ubiquitination was detected by immunoblotting with an anti-ubiquitin antibody (top panel) and unmodified Dma1 was detected with fluorescently-labelled streptavidin (bottom panel). (E) Relative protein levels of Dma1 (top) in the indicated strains as determined by immunoblotting immunoprecipitates relative to Cdc2 in the lysates (bottom) (upper panel) followed by quantification with Odyssey (lower panel). Quantification data is average ± standard deviation from two independent experiments. (F) Recombinant MBP-Dma1 was incubated with an E1-activating enzyme and/or the E2-conjugating enzyme, UbcH5a/UBE2D1, and methylated ubiquitin. Ubiquitin-modified Dma1 was detected by immunoblotting with an anti-ubiquitin antibody (top panel) and unmodified Dma1 was detected with anti-Dma1 serum (bottom panel). (G) Recombinant MBP-Dma1 proteins were incubated with an E1-activating enzyme, the E2-conjugating enzyme UbcH5a/UBE2D1 and methylated ubiquitin. Dma1 was cleaved from MBP and auto-ubiquitination was detected by immunoblotting with anti-Dma1 serum. (H) Recombinant MBP-Dma1 proteins were incubated with an E1-activating enzyme, the E2-conjugating enzyme ubcH5a/UBE2D1, and methylated ubiquitin. MBP-Dma1 ubiquitination was detected by immunoblotting with anti-Dma1 serum.

To determine if the different intensities measured at SPBs reflected increased abundance of Dma1-I194A relative to wildtype Dma1, we examined protein levels by whole cell fluorescence intensity and immunoblotting (Figure 2B and C). By both methods, the mutant protein was 2.6-fold higher in abundance, indicating that inactivating the catalytic activity of Dma1 causes an increase in total Dma1 protein. However, the overall difference in protein abundance was not as high as the difference in SPB intensity (2.6-fold increase vs. 3.2-fold increase) and so cannot fully account for the increased protein and lack of dynamics of Dma1-I194A observed at SPBs. Dma1 auto-ubiquitinates *in vitro* (Johnson et al., 2012a; Wang et al., 2012). Therefore, we next asked whether a lack of auto-ubiquitination in catalytically inactive Dma1-I194A could explain its increased SPB localization abundance and decreased SPB localization dynamics.

To establish that Dma1 auto-ubiquitinates *in vivo*, Dma1 and Dma1-I194A, both tagged with his_6_-biotin-his_6_ (HBH), were purified under fully denaturing conditions and probed for the presence of ubiquitinated forms by immunoblotting. Using the *mts3–1* proteasome mutant to block turn-over of ubiquitinated proteins, we detected ubiquitin modification of wildtype Dma1 but not catalytically inactive Dma1-I194A (Figure 2D) (Gordon et al., 1996). Thus, Dma1 auto-ubiquitinates *in vivo* as well as *in vitro*.

To test the idea that Dma1 auto-ubiquitination might promote its proteasomal degradation, we measured Dma1 levels in *mts3–1* relative to wildtype and found that Dma1 was more abundant in *mts3–1* arrested cells (Figure 2E). These data are consistent with auto-ubiquitination triggering Dma1 destruction.

To investigate directly if ubiquitin modifications affect Dma1’s ability to localize to SPBs, we sought to identify and mutate the auto-ubiquitinated residues. We first examined the sites of Dma1 auto-ubiquitination *in vitro* using recombinant Dma1 (Figure 2F) (Johnson et al., 2012a; Wang et al., 2012). Dma1 contains fourteen lysines that could be targeted for auto-ubiquitination. A series of mutants containing increasing numbers of lysine to arginine mutations was generated recombinantly and assayed for auto-ubiquitination using methyl ubiquitin to prevent chain elongation. Though the number of ubiquitin moieties added to Dma1 decreased with decreasing availability of lysines, only lysine-less Dma1 (Dma1–14KR) was completely unmodified (Figure 2G). Dma1 also ubiquitinated its MBP tag if the tag was not removed prior to the ubiquitination reaction (Figure 2H), indicating that Dma1 promiscuously ubiquitinates lysines in its proximity.

To determine which Dma1 lysines are ubiquitinated *in vivo*, we performed LC-MS/MS analyses on Dma1 purified from cells at different cell cycle stages. Nine ubiquitinated lysines of the 14 possible were identified (Figures S3A-I and S4A). We constructed mutant strains in which the 9 identified modified lysines or all 14 lysines in the protein were substituted with arginine (9KR and 14KR, respectively). Surprisingly, both Dma1 mutants tagged endogenously with the HBH tag were ubiquitinated (Figure S4B). The abundance of Dma1–14KR was significantly reduced, likely because so many mutations disrupted its structure (Figure S4B). However, that both were still ubiquitinated, combined with the ability of Dma1 and Dma1–14KR to ubiquitinate an MBP tag *in vitro* (Figure 2H), suggested that tags on the protein could be ubiquitinated in lieu of, or in addition to, Dma1 lysines. Indeed, LC-MS/MS analyses of Dma1-TAP and Dma1-HBH purified from cells identified ubiquitinated lysine residues in both tags (Figure S3J and K), confirming that Dma1 ubiquitinates its tags both *in vitro* and *in vivo*. Therefore, we predict that although the Dma1–9KR-mNG and Dma1–14KR mutants exhibited wildtype localization dynamics (Figure S4C and D, data not shown), this is because Dma1 is still able to ubiquitinate the mNG tag.

We next reasoned that wildtype Dma1 would be able to auto-ubiquitinate an inactive form of itself in close proximity and therefore restore wildtype dynamics to the catalytically inactive mutant. To test this, we constructed three diploid strains containing one mNG-tagged allele: *dma1-mNG/dma1^+^; dma1-I194A-mNG/dma1^+^; dma1-I194A-mNG/dma1-I194A*. When both alleles were wildtype or both inactive, the localization dynamics of the tagged protein mirrored the haploid situation, with Dma1-mNG dimming or disappearing from 100% of 20 SPBs during anaphase (Figure 3A and D) and Dma1-I194A-mNG showing no change at 84% of 44 SPBs (Figure 3B and E). In contrast, when Dma1-I194A-mNG was combined with wildtype Dma1, the tagged inactive protein now exhibited dynamics similar to wildtype, with the Dma1-I194A-mNG signal dimming or disappearing at 93% of 28 SPBs during anaphase (Figure 3C and F). This result indicates that the untagged wildtype Dma1 ubiquitinates the inactive Dma1-I194A-mNG, a reaction that could occur in *cis* or *trans* because Dma1 is a dimer (Johnson et al., 2012a). These results are consistent with a model in which auto-ubiquitination triggers the transient loss of Dma1 from SPBs during early anaphase B to allow SIN activation.

**Figure 3.**
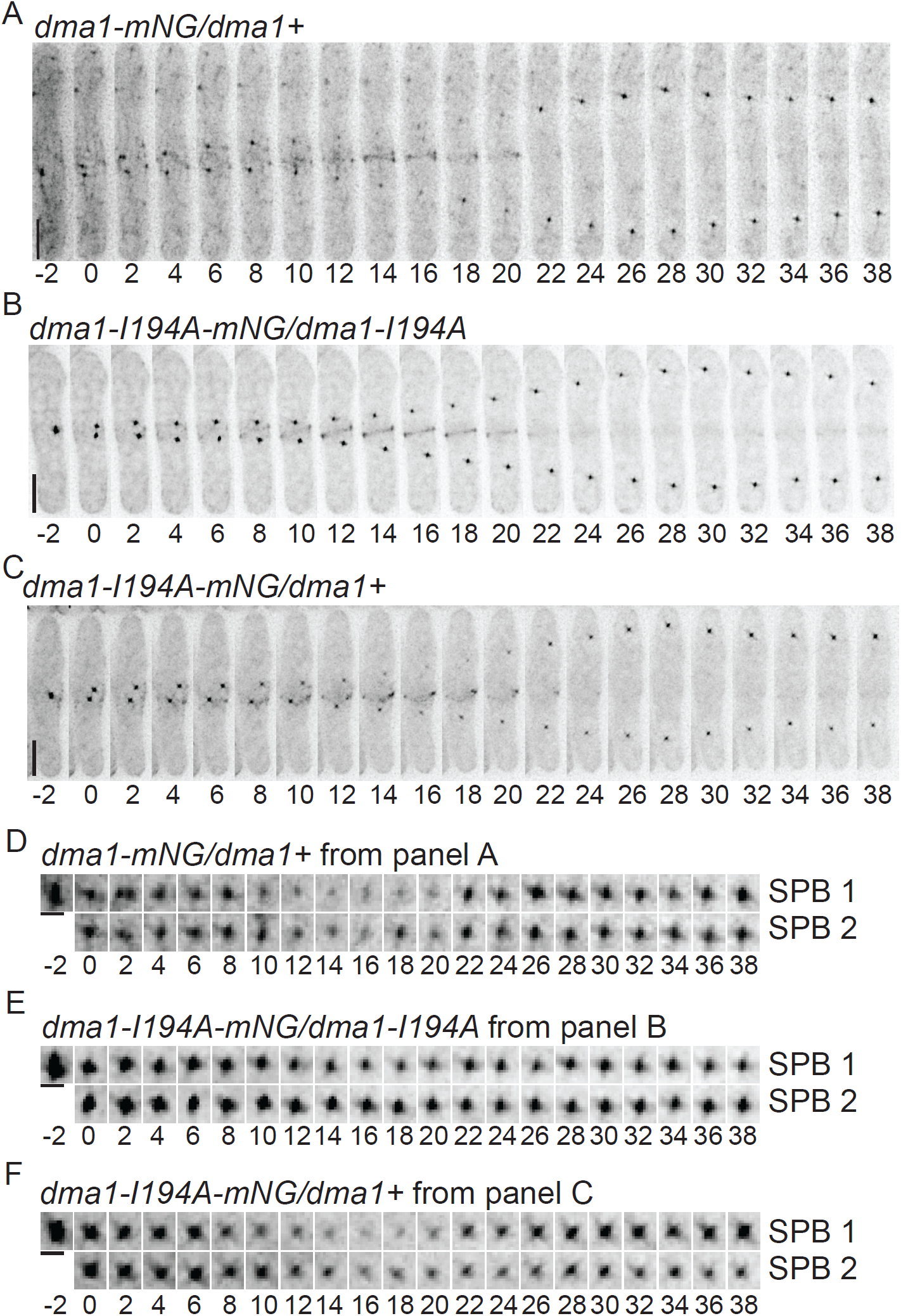
Dma1 catalytic activity drives its localization dynamics. (A-C) Images from representative movies of *dma1-mNG/dma1+* (A), *dma1-I194A-mNG/dma1-I194A* (B), and *dma1-I194A-mNG/dma1+* (C). Time in minutes denoted below images. Time 0 indicates initial frame of SPB separation. Scale bars, 5 μm. (D-F) Enlarged SPB region(s) from movies in A-C. Scale bars, 1 μm.

To test directly whether Dma1 auto-ubiquitination prevents its binding to phosphorylated Sid4, a step necessary for SIN inhibition, full-length Dma1 was purified from bacteria. Like the FHA domain alone (Johnson et al., 2013), full-length Dma1 preferentially bound phosphorylated Sid4 peptide (Figure 4A). When Dma1 was allowed to auto-ubiquitinate prior to the binding reaction, it did not bind the Sid4 phosphopeptide, and the ubiquitinated Dma1 was detected in the supernatant (Figures 4A and B). To clearly detect unbound ubiquitinated Dma1, the supernatants of the binding reaction were incubated with USP2, a commercially available deubiquitinating enzyme that shows activity towards multiple types of ubiquitin chains in vitro (Komander et al., 2009). USP2 treatment allowed detection of Dma1 in the supernatants and confirmed that it did not bind the beads (Figure 4A), confirming that Dma1 can only interact with Sid4 phosphopeptide if it is not ubiquitinated. If Dma1 was bound to the Sid4 phosphopeptide before the ubiquitination reaction, it was released into the supernatant (Figure 4C and D). These data support the idea that auto-ubiquitination at the onset of anaphase B triggers Dma1 removal from phosphorylated Sid4 at SPBs to allow SIN activation.

**Figure 4.**
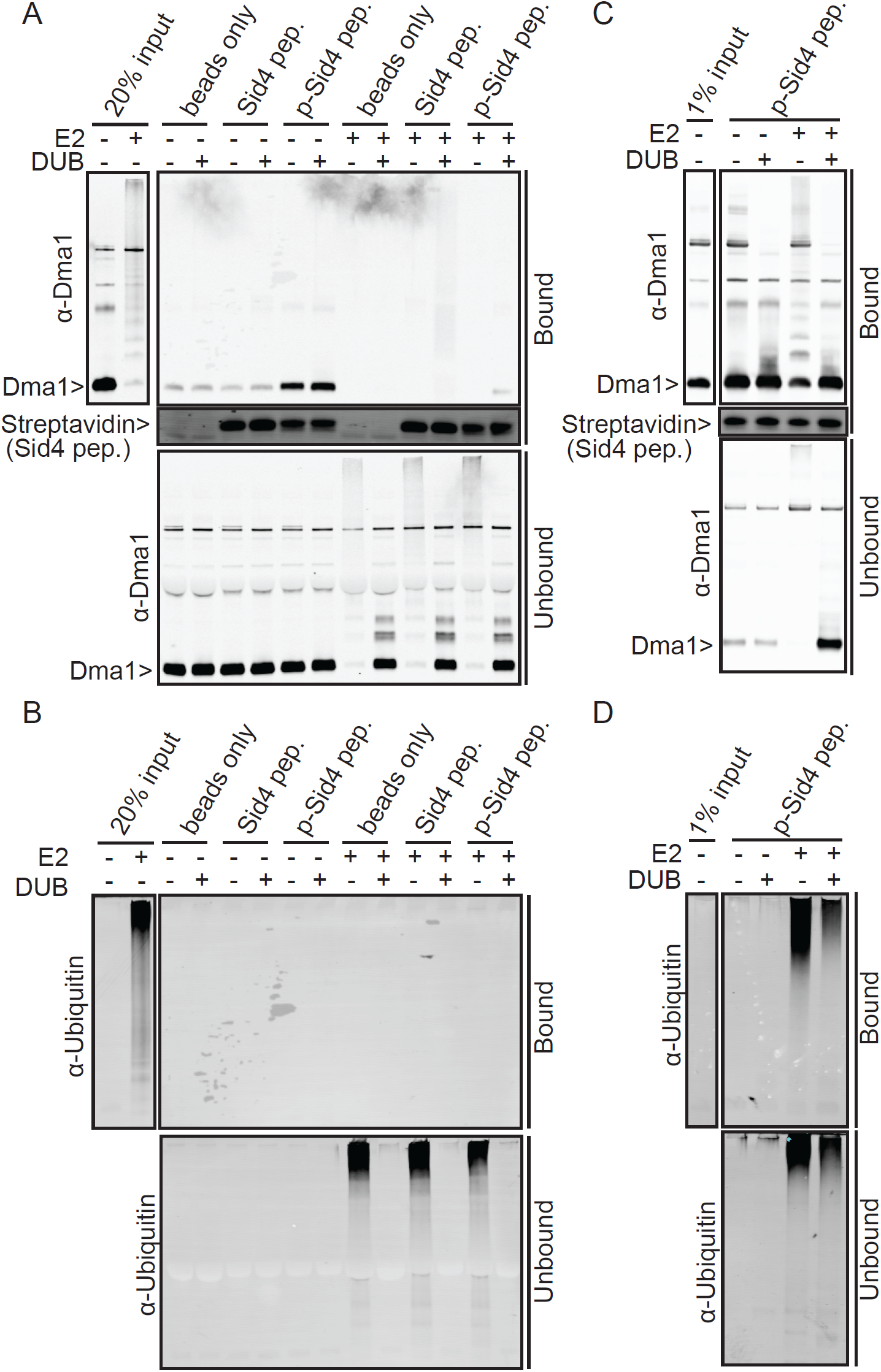
Dma1 must leave the SPBs for cytokinesis. (A) Recombinant Dma1 was incubated with E1, Ub and E2 (+E2) or not (−E2), followed by incubation with biotinylated Sid4 peptides (either unphosphorylated (Sid4) or phosphorylated (p-Sid4)) conjugated to streptavidin beads. Beads only was included as a control. Supernatants (containing unbound Dma1) and beads (containing bound Dma1) were recovered and each was split in half for treatment either with H_2_O, or USP2 to collapse ubiquitinated Dma1 for better detection. The entire samples from beads (upper panel) and half of the supernatants (lower panel) were resolved by SDS-PAGE and detected by immunoblotting with anti-Dma1 serum or streptavidin (for visualization of biotinylated peptides). (B) anti-Ub blotting of the same membranes blotted for anti-Dma1 in Figure 4A. (C) Recombinant Dma1 was incubated with biotinylated phosphorylated Sid4 (p-Sid4) peptides. The beads were split in half and subjected to *in vitro* ubiquitination assay with (+E2) or without (−E2) E2 enzyme. Supernatants (containing unbound Dma1) and beads (containing bound Dma1) were recovered and each was split in half for treatment either with H_2_O or USP2. The entire samples from beads (upper panel) and supernatants (lower panel) were resolved by SDS-PAGE and detected by immunoblotting with anti-Dma1 serum or streptavidin. (D) anti-Ub blotting of the same membranes blotted for anti-Dma1 in Figure 4C.

## Relationship between Dma1 SPB dynamics and the SIN

The dynamic localization of Dma1 during mitosis is reminiscent of that displayed by several SIN components (Johnson et al., 2012b; Simanis, 2015). Asymmetric SPB localization of Cdc7 to one SPB and localization of Sid2 to the division site are considered markers of maximal SIN activation and Cdk1 inhibition (Johnson et al., 2012b; Simanis, 2015). Therefore, we imaged Dma1-mNG in combination with those two SIN components (Figure 5) to place the timing of dynamic Dma1 localization in the context of SIN activation. Dma1-mNG SPB dimming occurred 1–2 minutes in 14 cells and 3–5 minutes in 3 cells prior to the development of detectable asymmetry in Cdc7-mCherry signal in the 17 cells examined (Figure 5A and B), and the transient reduction in Dma1 division site localization preceded Sid2 division site localization in all 8 cells examined (Figure 5C).

**Figure 5.**
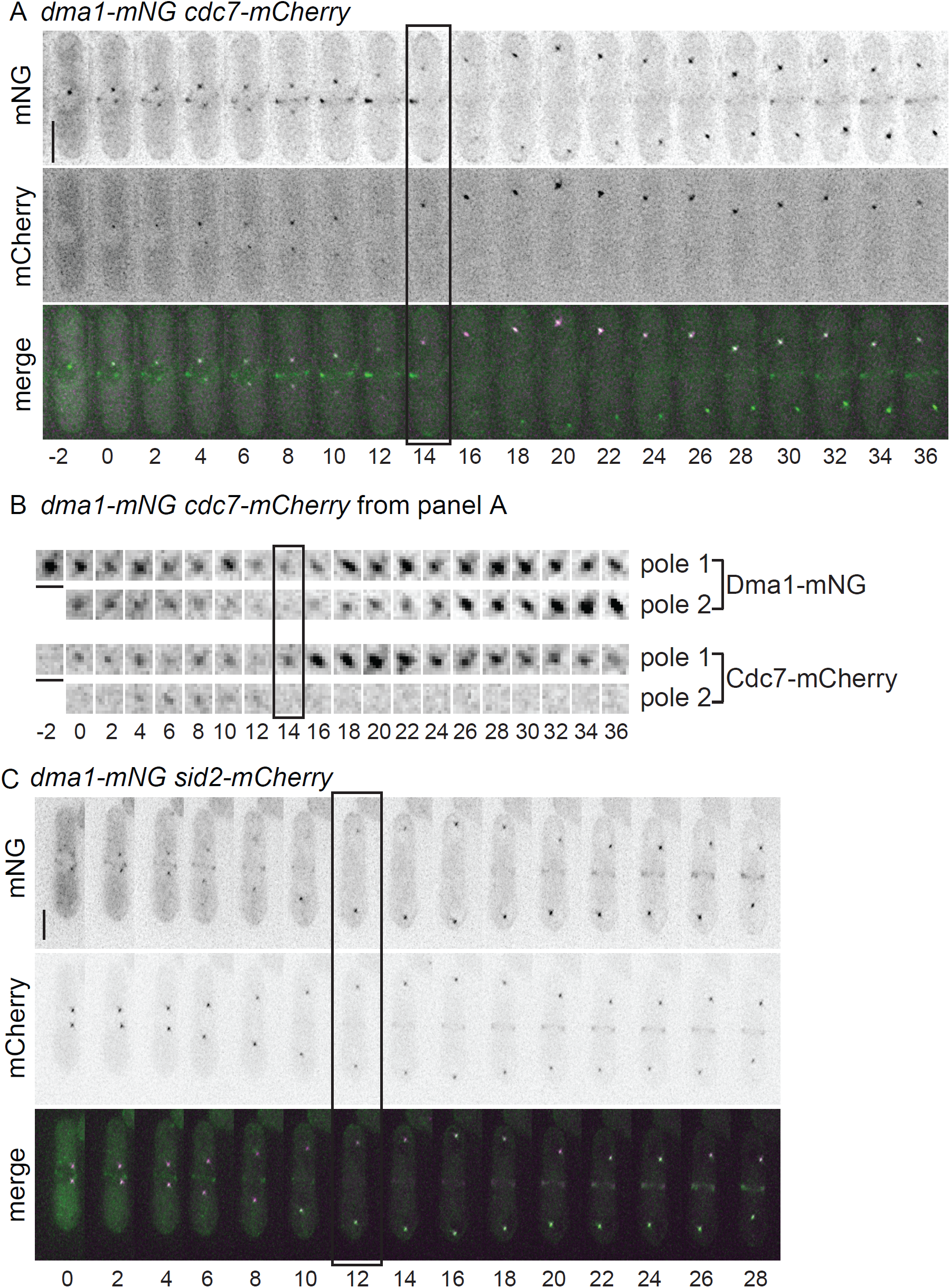
SPB dynamics of Dma1 relative to SIN components. Images from representative movies of *dma1-mNG cdc7-mCherry* (A) and *dma1-mNG sid2-mCherry* (C). Time in minutes denoted below images. In A, time 0 indicates initial frame of SPB separation while in C, time 0 indicates the beginning of imaging. Scale bars, 5 μm. (B) Enlarged SPB region(s) from movie in A. Scale bars, 1 μm. In A and B, boxes indicate initial frame of asymmetric Cdc7 SPB localization. In C, the box indicates the first frame Sid2 localization to the division site was detected.

To test whether SIN activity modulated some aspect of Dma1’s localization pattern, Dma1-mNG was imaged in the SIN mutant *cdc7–24* as cells passed through mitosis. Dma1-mNG intensity dimmed or disappeared at 21 of 22 SPBs in 11 cells but it never re-intensified at SPBs as in wildtype cells (Figure 6A and B), indicating that SIN activity is important for Dma1 re-accumulation at SPBs in late anaphase. Dma1 did re-intensify at the division site in all 11 cells examined. We next imaged Dma1-mNG in the hyperactive SIN mutant *cdc16–116*. In these cells that have constitutive SIN activity (Fankhauser et al., 1993), Dma1-mNG was detected on 97% (116/119) of SPBs (Figure 6C). Since in wildtype cells the SIN is not maximally active until Cdk1 activity falls later in anaphase (He et al., 1997; Guertin et al., 2000; Dischinger et al., 2008), these results raise the possibility that high SIN activity promotes Dma1 SPB re-accumulation at the end of anaphase.

**Figure 6.**
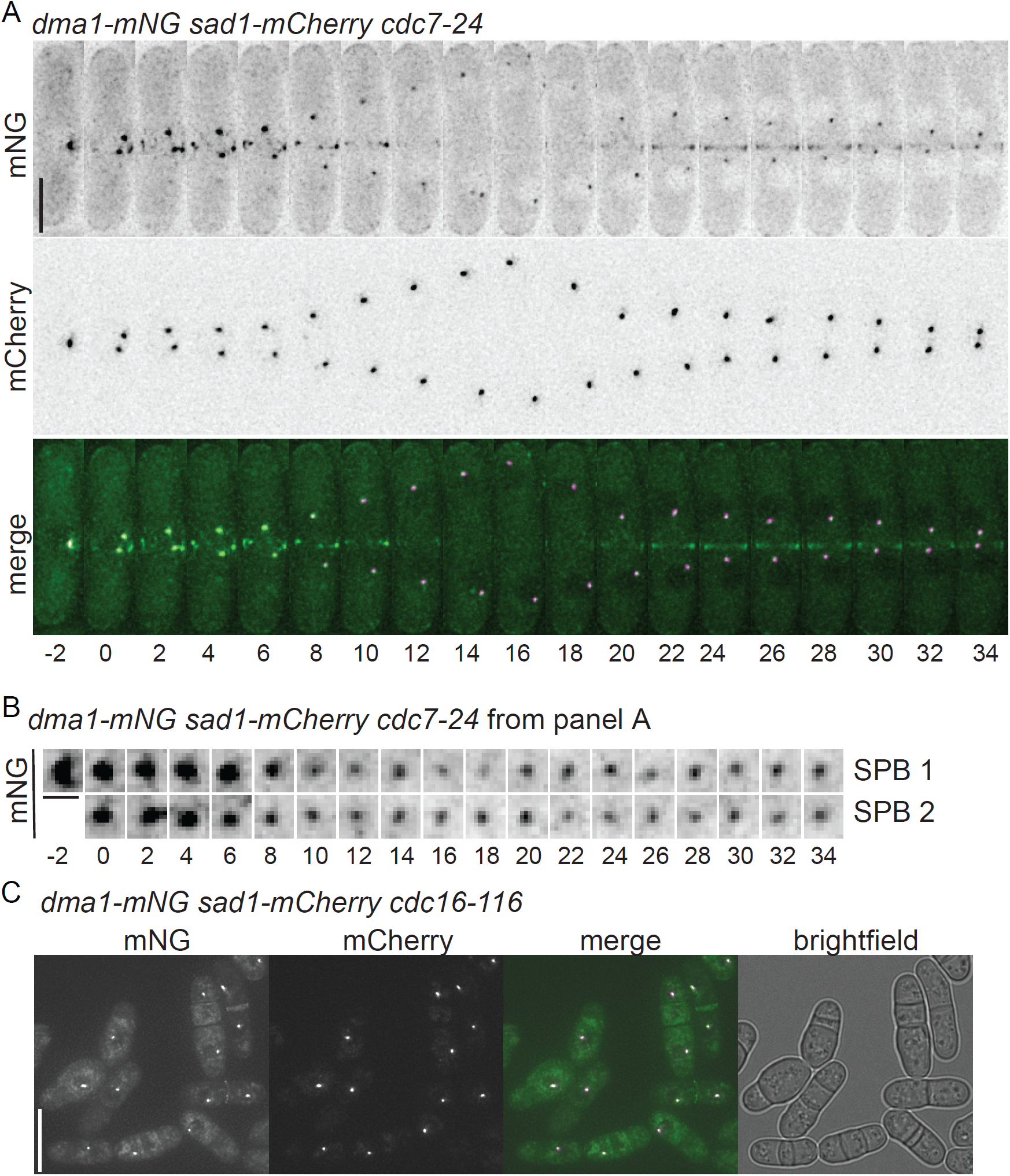
Influence of SIN function on Dma1 localization dynamics. (A) Images from a representative movie of *dma1-mNG sad1-mCherry cdc7–24* following shift to 36°C for 2 h. Time in minutes denoted below images; time 0 indicates initial frame of SPB separation. Scale bar, 5 μm. (B) Enlarged SPB region(s) from movie in A. Scale bars, 1 μm. (C) Representative live cell image of *dma1-mNG sad1-mCherry cdc16–116* shifted to 36°C for 3 h. Scale bar, 10 μm.

## Constitutive Dma1 localization to SPBs prevents cytokinesis

We next wanted to determine the importance of Dma1 transiently cycling off of SPBs during anaphase B by permanently tethering it to SPBs. To do this, we tagged Sid4 at its endogenous locus with GFP-binding protein (GBP), which has a high affinity for GFP (Rothbauer et al., 2006; Rothbauer et al., 2008). When a *sid4-GBP-mCherry* or *sid4-GBP* strain was crossed to *dma1-GFP*, many double mutants (31/35 and 7/19, respectively) were dead. Similarly, synthetic lethality was previously observed when the tags were reversed and *dma1-GBP-mCherry* was expressed in a *sid4-GFP* strain (Chen et al., 2017), though in this study, co-localization at SPBs and dependency of cell death on failed cytokinesis was not determined. Fortunately, we were able to recover some double-tagged strains for imaging, although they grew very poorly: 22% of *dma1-GFP sid4-GBP-mCherry* and 11% of *dma1-GFP sid4-GBP* cells were dead or lysing. In live *dma1-GFP sid4-GBP-mCherry* cells, Dma1-GFP localized to the division site normally and co-localized with Sid4-GBP-mCherry at SPBs at all stages of the cell cycle (Figure 7A), indicating that Dma1-GFP was tethered permanently to SPBs. In both double mutant strains, there was a higher percentage of multi-nucleated cells than in wildtype and 46% of bi-nucleated cells had “kissing” nuclei (Figure 7B-D) indicative of SIN failure (Hagan and Hyams, 1988). We reasoned that if the SIN was inhibited in these strains via Dma1-mediated Sid4 ubiquitination, introducing the *ppc89-DUB* fusion allele that prevents Sid4 ubiquitination (Figure S2B) should rescue growth of *dma1-GFP sid4-GBP-mCherry* cells. As predicted, of the 14 *dma1-GFP sid4-GBP-mCherry ppc89-DUB* triple mutant strains constructed, all were viable and showed reduced levels of multinucleation and “kissing” nuclei (Figure 4D). These data indicate that constitutive association of Dma1 with Sid4 drives Sid4 ubiquitination, compromises SIN activity, and results in cytokinesis failure. Thus, Dma1’s transient loss from SPBs at the onset of anaphase must be a critical step in silencing the mitotic checkpoint, as well as allowing cytokinesis to proceed in unperturbed cells.

**Figure 7.**
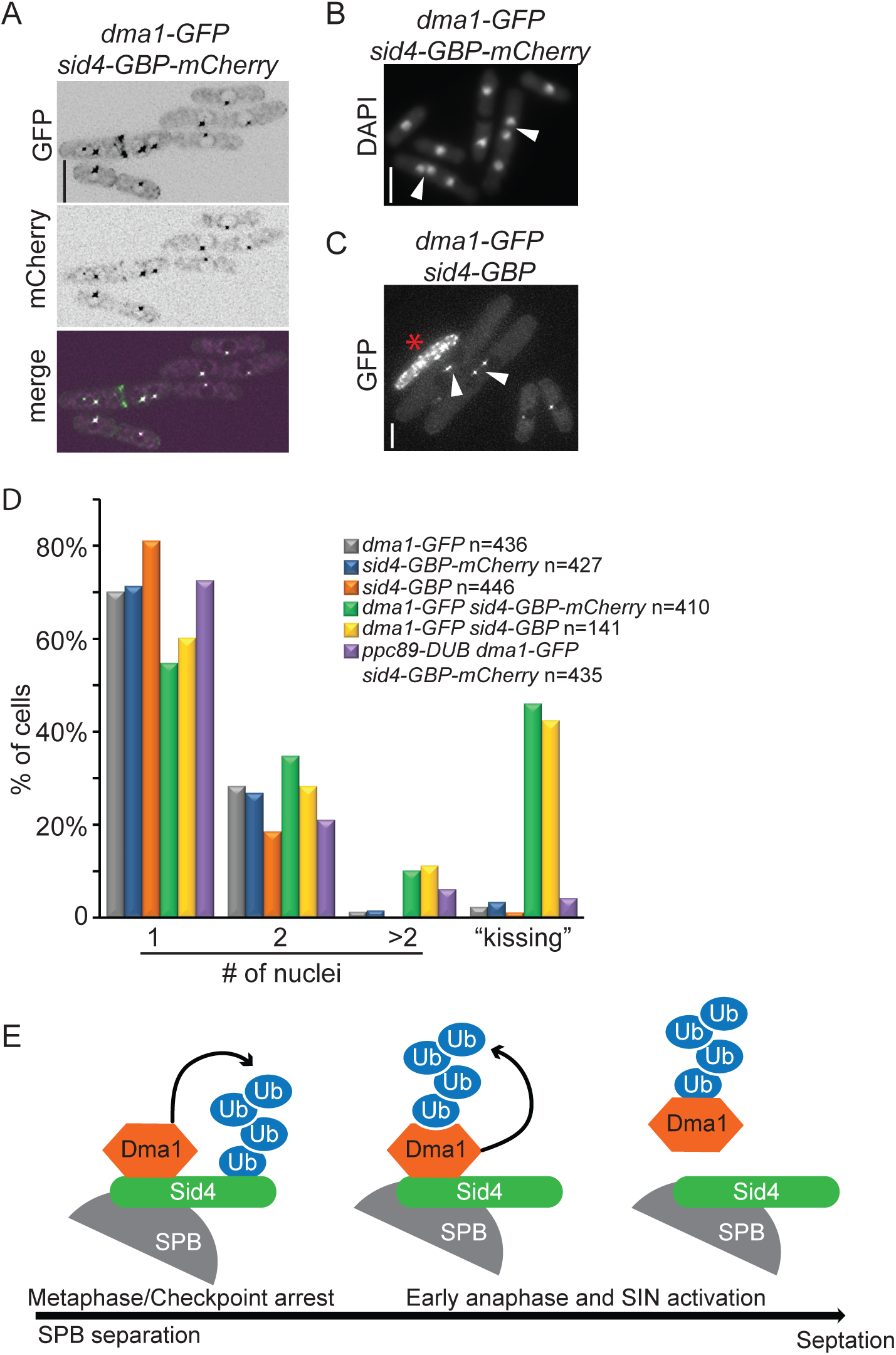
Dma1 must leave the SPBs for cytokinesis. (A) Representative live cell images of *dma1-GFP sid4-GBP-mCherry*. (B) Representative DAPI stained images of *dma1-GFP sid4-GBP-mCherry*. (C) Representative image of *dma1-GFP sid4-GBP* cells illustrating “kissing nuclei” phenotype indicated with arrows in B and C. (D) Quantification of cells from B and C with 1, 2, and >2 nuclei and separately “kissing nuclei”. (E) Cartoon showing the dynamic localization of Dma1 throughout mitosis and cytokinesis. In metaphase when the mitotic checkpoint is activated, Dma1 binds and ubiquitinates Sid4. Then, in early anaphase, Dma1 auto-ubiquitinates and leaves the SPB allowing maximal SIN activation and cytokinesis.

## Discussion

SPB localization of the ubiquitin ligase Dma1 is required for its function in a mitotic checkpoint that stalls cytokinesis through SIN inhibition when a mitotic spindle cannot form (Murone and Simanis, 1996; Guertin et al., 2002). In this report, we show that Dma1 exhibits previously unrecognized dynamic localization to SPBs and the cell division site during anaphase. We found that both the SIN and auto-ubiquitination modulate Dma1 SPB localization dynamics, and therefore its function in the checkpoint. Further, our data show that if Dma1 is prevented from leaving Sid4 on the SPBs, cells fail division, indicating that Dma1’s transient loss from SPBs is a critical step in cytokinesis and cell survival.

### Dma1 division site localization

Though Dma1 was previously shown to localize to the division site (Guertin et al., 2002), our time-lapse imaging experiments with the brighter mNG fluorophore (Shaner et al., 2013) revealed more detail. Dma1-mNG initially appears in node-like structures, reminiscent of cytokinetic precursor nodes (Rincon and Paoletti, 2012), before coalescing into a ring. In addition, Dma1 leaves the division site after anaphase B onset and before ring constriction and then re-appears at the division site before cell division. The role of Dma1 at the division site is not yet understood because no binding partners or substrates at the medial cortex have been identified. Because neither Dma1 activity nor the SIN impacts its cell division site localization dynamics, a different mechanism of targeting to this site must be in place. Importantly, tethering Dma1 to Sid4 at the SPB using the GBP-GFP system blocked cytokinesis without disrupting Dma1 localization to the division site, validating that it is Dma1’s SPB localization that is important for controlling SIN function. We also detected Dma1 localization to cell tips in this study, but again, its mechanism of targeting and function there are not known.

### SIN regulation of Dma1 dynamics

Although the transient dip in Dma1 SPB localization in early anaphase depends on its catalytic activity and is independent of SIN function, SIN activity is required for Dma1’s re-accumulation at SPBs later in anaphase. This observation may reflect a change in Dma1’s auto-ubiquitination activity or that of a counter-acting deubiquitinating enzyme, or reveals a component of negative feedback in which the SIN activates an inhibitor to direct its own silencing (Garcia-Cortes and McCollum, 2009). Since Dma1 is a phosphoprotein (Koch et al., 2011), it will be interesting to determine the role of phosphorylation in Dma1 regulation and whether this is controlled directly or indirectly by SIN kinases.

### Dma1 auto-ubiquitination controls its dynamic localization at SPBs

Our data show that Dma1 is capable of extensive auto-ubiquitination *in vitro* and *in vivo*. Its promiscuous auto-ubiquitination, which extended to any tag that we tested that includes lysines, complicated our investigation of its role as we were unable to visualize the dynamic localization of a lysine-less catalytically-active Dma1 protein with confidence in a haploid cell. However, we were able to use heterozygous diploids to show that the active form of the protein was sufficient to remove the inactive form from SPBs during early anaphase. Indeed, our data support the idea that auto-ubiquitination at the onset of anaphase triggers transient Dma1 removal from SPBs to allow full SIN activation (Figure 7E). By preventing Dma1 from leaving the SPB using the GBP trap, the SIN was inhibited. These results are consistent with the role of Dma1 as a SPB-localized SIN antagonist in early mitosis whose function must be relieved at the onset of anaphase (Guertin et al., 2002; Johnson and Gould, 2011; Johnson et al., 2013).

### Degradation of Dma1

Because the catalytically inactive form of Dma1 is more abundant than wildtype, its levels are clearly regulated by its own activity state. While we do not yet know all the factors modulating Dma1 catalytic activity, our data is consistent with auto-ubiquitination triggering Dma1 destruction. This could happen directly at SPBs, analogously to proteasome-mediated degradation of many regulatory proteins at centrosomes (Vora and Phillips, 2016), or off the centrosome if ubiquitination prevents Dma1 association with its SPB tethers, similar to what has been observed for another SIN inhibitor, Byr4 (Krapp et al., 2008). This mechanism of inhibiting Dma1, auto-ubiquitination and self-destruction, may be broadly applicable in signaling pathways when the outcome of substrate ubiquitination is not degradation and signal flux through a pathway must be restored rapidly. Dma1 relatives CHFR and RNF8 are integral components of checkpoint pathways that undergo auto-ubiquitination *in vivo* and *in vitro* (Kang et al., 2002; Lok et al., 2012), but whether this modification is a key step in checkpoint silencing in their cases remains to be determined. To fully understand the factors that modulate the Dma1-dependent mitotic checkpoint, it will be important to determine how Dma1 auto-ubiquitination is restrained to allow substrate ubiquitination during early mitosis. It will also be important to understand the checkpoint silencing strategies used in other organisms to allow cytokinesis to proceed following chromosome clearance from the division site and whether there are parallels with the strategies used by fission yeast.

## Materials and Methods

### Yeast

Yeast strains grown in this study (Table S1) were grown in yeast extract (YE), 4X concentrated yeast extract (4X YE) or minimal media supplemented with leucine and uracil for diploid strains (Moreno et al., 1991). Genes were tagged at the 3’ end of their open reading frame (ORF) with *mNG:kan^R^, mNG:Hyg^R^, GFP:kan^R^, mCherry:kan^R^*, *mCherry:nar, HBH:kan^R^, HA_3_-TAP:kan^R^, GBP-mCherry:kan^R^*, or *ubp7^+^* (residues 201–875) *DUB:kan^R^* using PCR and pFA6 cassettes (Wach et al., 1994; Bahler et al., 1998). For gene replacements, a haploid *dma1::ura4^+^* strain was transformed with the appropriate pIRT2–*dma*1 mutant, integrants were selected by resistance to 1.5 mg/mL 5-FOA, and validated by colony PCR and DNA sequencing. Diploid strains were made by crossing *ade6-M210* cells with *ade6-M216* cells on glutamate plates for 24–48 h followed by re-streaking to single colonies on MAU plate and incubating at 32°C for 3 days. White colonies (diploid cells) were then picked for subsequent analysis. For spot assays, cells were cultured in YE at 29°C. Serial 10-fold dilutions of each culture were made, 3 μL of each dilution was spotted on YE plates and they were incubated at various temperatures for 3–4 days. *nda3-KM311* cultures were grown at 32°C and then shifted to 18°C for 6 h to block in prometaphase. *mts3–1* cells were grown at 25°C and then shifted to 36°C for 3.5 h before imaging or biochemical experiments.

### Molecular biology

Plasmids were generated by standard molecular biology techniques. *dma1* mutations were made either in the context of a gene fragment in the pIRT2 vector that included 500 bp upstream and downstream of the open reading frame or in the context of the open reading frame in pMAL-c2 vector using a QuikChange site-directed mutagenesis kit (Agilent Technologies). The K to R mutation abbreviations are as follows: <u>Dma1–4KR</u> = K3R, K10R, K121R, and K124R; <u>Dma1–9KR</u> = K3R, K26R, K54R, K82R, K124R, K164R, K174R, K237R, and K262R; <u>Dma1–12KR</u> = Dma1–4KR mutations plus K22R, K26R, K162R, K164R, K174R, K217R, K237R, and K262R; <u>Dma1–14KR</u> = Dma1–12KR mutations plus K54R, and K82R.

cDNAs encoding Ubp2, Ubp3, Ubp4, Ubp8, Ubp14, Ubp15, Ubp16, Uch1, were amplified by PCR from genomic *S. pombe* DNA using primers containing restriction sites. PCR products were digested with restriction enzymes (Nde1/BamHI for *ubp2* and *ubp3* and NdeI/XmaI for *ubp4, ubp8, ubp14, ubp15, ubp16*, and *uch1*), subcloned into pREP1, and verified by sequencing. DNAs encoding Ubp1, Ubp7, Ubp9, Ubp11, Otu1, Otu2, and Sst2, were digested from genomic constructs, subcloned into pREP1 and verified by sequencing. *ubp6* and *uch2* cDNAs in pREP1 vectors were gifts from Dr. Colin Gordon (Stone et al., 2004).

### Microscopy

Unless otherwise indicated, all imaging was done at room temperature on a Personal DeltaVision microscope system (Applied Precision) that includes an Olympus IX71 microscope, 60× NA 1.42 PlanApo oil immersion objective, standard and live-cell filter wheels, a Photometrics CoolSnap HQ2 camera, and softWoRx imaging software. Time-lapse imaging was done using a CellAsic microfluidic system (EMD Millipore) with appropriate cell media. Figure images are maximum intensity projections of z sections spaced 0.2–0.5 μm apart. Images used for quantification were not deconvolved and sum projected.

Intensity measurements were made with ImageJ software (National Institutes of Health, Bethesda, MD: http://rsbweb.nih.gov/ij/). For all intensity measurements, the background was subtracted by calculating a value for background per pixel (BPP) in the same image as the region of interest (ROI) where there were no cells. The BPP was then multiplied by the area of the ROI and the resulting product was subtracted from the raw integrated intensity of that ROI yielding the corrected intensity value (Waters, 2009).

For analysis relative to a SPB marker, an ROI was made based on the SPB marker. mNG or GFP fluorescence intensity and mCherry marker fluorescence intensity were measured with background correction for each. Final values for each cell are expressed as mNG/mCherry or GFP/mCherry ratios. Measurements for the cells in each group were averaged for statistical analysis.

### Protein purification and mass spectrometry

Endogenously tagged versions of Dma1 (Dma1-HA_3_-TAP and Dma1-HBH) were purified as previously described (Gould et al., 2004; Tagwerker et al., 2006; Elmore et al., 2014) and analyzed by 2D-LC-MS/MS as previously described (McDonald et al., 2002; Roberts-Galbraith et al., 2009). RAW files were processed using two pipelines: 1) using Myrimatch (v 2.1.132) (Tabb et al., 2007) and IDPicker (v 2.6.271.0) (Ma et al., 2009) as previously described (McLean et al., 2011) and 2) using turboSEQUEST, Scaffold (v 4.4.7) and Scaffold PTM (v 3.0.0) as previously described (Beckley et al., 2015). We detected the variable di-Gly modification (114 Da) indicating ubiquitination, and these modifications were manually validated (representative spectra are shown in Figure S3).

### Protein expression and purification

*dma1* variants were cloned into pMAL-c2 or pET-His_6-_MBP prescission LIC cloning vector (HMPKS) (HMPKS was a gift from Scott Gradia (Addgene plasmid # 29721)) for production as maltose-binding protein (MBP) fusions. Proteins were induced in *Escherichia coli* Rosetta2(DE3)pLysS cells by addition of 0.8 mM IPTG and overnight incubation at 18°C. Cells were lysed using 300 ug/mL lysozyme for 20 minutes followed by sonication. Proteins were affinity purified on pre-washed amylose resin (New England Biolabs) in MBP column buffer (100 mM NaCl, 20 mM Tris-HCl pH 7.4, mM EDTA, 1 mM DTT, and 1% Nonidet P40). Protein was eluted with 10 mM maltose and, in some cases, the MBP tag was cleaved with Factor Xa protease (New England Biolabs) or Precission protease (GE Healthcare). Dma1 peak elution fraction(s) were detected by SDS PAGE and Coomassie Blue (Sigma) staining. After pooling peak fractions, Dma1 proteins were concentrated using a 10 MWCO Amicon Ultra centrifugal filter (EMD Millipore). Dma1 was cleaved from MBP by adding 1 ul (2 units) of PreScission protease to 500 ug MBP-Dma1 on amylose beads (in 200 ul MBP column buffer) and incubating at 4°C overnight. The supernatant containing cleaved Dma1 was retrieved and the concentration of Dma1 was determined by SDS-PAGE using BSA as standard.

### *In vitro* ubiquitination assay

Ubiquitination reactions included: 0.25 to 1 μg recombinant MBP-Dma1 (or variant thereof), or Dma1 cleaved from MBP, 175 nM E1 (R&D Systems), 3 μM E2 (R&D Systems), 50 μg/mL methylated ubiquitin, 5 mM ATP, and 1x ubiquitination buffer (50 mM Tris-HCl pH 7.5, 2.5 mM MgCl_2_, 0.5 mM DTT). These 20 μL reactions were incubated with agitation at 30°C for 90 min before adding SDS sample buffer or cleaving Dma1 from MBP. For MBP cleavage, 1.8 μL of 1M CaCl_2_ and 5 μL Factor Xa were added and the reactions were incubated at room temperature with agitation for 1 h. Then, 100 μL benzamidine beads (1:1 slurry) and 100 μL amylose beads (1:1 slurry) (both pre-washed in MBP column buffer) were added and reactions were incubated room temperature with agitation for 45 min. Supernatants were transferred to new tubes and SDS sample buffer was added prior to separation by SDS-PAGE. Ubiquitination products and Dma1 were detected by immunoblotting with anti-ubiquitin (EMD Millipore, MAB1510) at a 1:250 dilution or (Life Sensors, VU-1) at a 1:500 dilution and/or anti-Dma1 (see Figure S4E) antibodies.

### *In vivo* ubiquitination assay

Dma1-HBH was purified from 250 mL 4X YE pellets as previously described (Tagwerker et al., 2006; Beckley et al., 2015). Briefly, cell pellets were washed with 10 mL modified buffer 1 (8 M urea, 300 mM NaCl, 50 mM NaPO_4_, 0.5% Nonidet P40 and 4 mM Imidazole, pH 8 with 1.3 mM benzamidine, 1 mM PMSF, 50 uM PR-619 (LifeSensors), 50 μM N-Ethylmaleimide, and 1 Complete Protease Inhibitor Cocktail Tablet, EDTA-free (Roche) per 50 mL), and then lysed by bead disruption with 500 μL buffer 1. Lysates were extracted with 10 mL then 5 mL buffer 1 (15 mL total extraction), cleared by high speed centrifugation, and then incubated with 200 μl (1:1 slurry) Ni^2^+-NTA agarose beads (Qiagen) for 3–4 h at room temperature. After incubation, beads were washed four times: 1X with 10 mL buffer 1 and 3X with 10 mL buffer 3 (8 M urea, 300 mM NaCl, 50 mM NaPO_4_, 0.5% Nonidet P40 and 20 mM Imidazole, pH 6.3). Proteins were then eluted for 15 min in 5 mL buffer 4 (8 M urea, 200 mM NaCl, 50 mM NaPO_4_, 0.5% Nonidet P40 and 2% SDS, 100 mM Tris and 10 mM EDTA, pH 4.3) 2X and the two eluates were pooled. The pH of the final eluate was adjusted to 8 before 80 μL streptavidin ultra-link resin (Pierce) was added and incubated overnight at room temperature. The streptavidin beads were then washed 3X with buffer 6 (8 M urea, 200 mM NaCl, 2% SDS and 100 mM Tris, pH 8). Purified proteins were detected on a western blot using an anti-ubiquitin antibody (EMD Millipore, MAB1510) at a 1:250 dilution or (Life Sensors, VU-1) at a 1:500 dilution and fluorescently labelled streptavidin (Li-COR Biosciences).

### Biotinylated peptide conjugation to streptavidin beads

20 ul of 50% slurry of ultralink strepavidin beads was washed in 200 ul MBP column buffer three times and resuspended in 88 ul column buffer. 2 ug (2 ul of 1 mg/ml) synthetic biotinylated peptides (Genescript) dissolved in 5% acetonitrile was added and incubated at 4°C overnight. Beads were then washed in 200 ul MBP column buffer three times before being resuspended in 10 ul column buffer.

### Sid4 phosphopeptide binding assay

The Sid4 phosphopeptide binding assay was performed as described (Johnson et al., 2013) with the following modifications. First, 0.5 ug purified recombinant Dma1 (cleaved from MBP-Dma1 by Precission protease) was ubiquitinated *in vitro* as described above (0.4 ug/ul ubiquitin was used instead of methylated ubiqutin) in the presence (+E2) or absence (−E2) of E2 enzyme in 40 ul total volume for 2 hours at 30°C. Reactions were stopped by adding 4 ul 1M DTT to a final concentration of 91.4 mM. Next, 2.2 ul of the reaction mixture containing 25 ng nonubiquitinated (−E2) or ubiquitinated (+E2) Dma1 was added to 19.8 ul binding buffer (50 mM Tris-HCl, pH 7.4, 25 mM KCl, 5 mM MgCl_2_, 1 mM DTT) containing streptavidin beads only, or 5 uM synthetic biotinylated nonphosphorylated Sid4 peptide (bio-LTSSTCVSSISQ) or phosphorylated Sid4 peptide (bio-LTSSpT_275_CVpS_278_SISQ) (Genescript) conjugated on ultralink streptavidin beads (Pierce) and incubated at 4°C for 30 min (beads were pre-blocked with 50 ul 5% BSA in binding buffer for 1h). The unbound supernatants were recovered, split equally into two parts, and treated with 0.33 ul H_2_O or 1.5 mg/ml USP2 (Boston Biochem) and analyzed by SDS-PAGE. The beads were washed with 1 ml high salt MBP column buffer (20 mM Tris-HCl, pH 7.4, 250 mM NaCl, 1 mM EDTA, 0.1% NP-40, 1 mM DTT) five times, followed by washing with 0.5 ml de-ubiquitination buffer (50 mM Tris, pH 7.4, 25 mM KCl, 5 mM MgCl_2_, 10 mM DTT) twice prior to being split equally into two parts. Beads were resuspended in 19 ul de-ubiquitination buffer and treated with 0.33 ul H_2_O or USP2. The de-ubiquitination reactions were incubated with shaking at 32°C for 2 h. Reactions were stopped by adding SDS sample buffer.

Alternatively, phosphorylated Sid4 peptides on beads (6.25 μM) were incubated with 5 ng/μL Dma1 in 200 μL high salt MBP buffer for 30 min at 4°C in the presence of 5% BSA, followed by 6 × 1 mL wash of high salt MBP buffer and then with 1 mL of ubiquitination buffer twice. Beads were split into two equal parts (for −E2 and +E2) and subject to the *in vitro* ubiquitination assay in 20 μL total volume as described above. Reactions were stopped by adding 0.5 uL 1M DTT to a final concentration of 24.9 mM. The unbound supernatants were recovered, split equally into two parts, and 10 μL of 50 mM Tris, 50 mM KCl 7.5 mM MgCl_2_ was added (after adjusting the buffer composition is: 49.4 mM Tris, 25 mM KCl, 5 mM MgCl_2_, 12.4 mM DTT) prior to being treated with 0.33 ul H_2_O or 1.5 mg/ml USP2. The beads were washed with 1 mL deubiquitination buffer twice and split equally into two parts before treatment with H_2_O or USP2 as described above.

### Lysis, immunoprecipitation, and immunoblotting

Cell pellets were washed once in NP-40 buffer (10 mM sodium phosphate pH 7.0, 1% Nonidet P40, 150 mM NaCl, 2 mM EDTA) with inhibitors (1.3 mM benzamidine, 1 mM PMSF, and 1 Complete Protease Inhibitor Cocktail Tablet, EDTA-free (Roche) per 50 mL) and lysed by bead disruption. For denaturing lysis, 500 μL SDS lysis buffer (10 mM sodium phosphate pH 7.0, 1 % SDS, 1 mM DTT, 1 mM EDTA, 50 mM NaF, 100 μM sodium orthovanadate, 1 mM PMSF, and 4 μg/mL leupeptin) was added and sample was incubated at 95°C for 2 min, lysate was extracted with 800 μL NP-40 buffer and transferred to a clean tube. For native lysis, the lysate was extracted with 500 μL NP-40 buffer and again with 800 μL, then transferred to a clean tube. Extractions were followed by two clearing spins of 5 and 30 min. Proteins were immunoprecipitated from native protein lysates using an excess of antibody (listed below) and rocking at 4°C for 1 h, followed by addition of Protein A or G Sepharose beads (GE Healthcare), as appropriate, and rocking at 4°C for 30 min. Samples were washed 4X with NP-40 buffer. Antibodies: 0.8 μg anti-GFP (Roche), 2 μg anti-FLAG (Sigma-Aldrich), 3 μL rabbit anti-Sid4 antiserum or 2 μL rabbit anti-Dma1 serum.

Antisera were raised against recombinant GST-mNG or GST-Dma1 (Cocalico) and their specificity was verified by immunoblotting (Figures 2 and S4E, respectively). The anti-Dma1 serum was further purified by ammonium sulfate precipitation. The serum was cleared by centrifugation and then precipitated with 0.5 volumes of saturated ammonium sulfate added dropwise and incubated overnight at 4°C. The precipitate was cleared from the serum by centrifugation then the serum was precipitated with an additional 0.5 volumes of saturated ammonium sulfate added dropwise and incubated overnight at 4°C. The precipitate was pelleted by centrifugation, resuspended in 0.4 volumes PBS, and dialyzed 3X in PBS.

For lambda phosphatase treatment, immunoprecipitated protein was washed twice with 25 mM HEPES-NaOH (pH 7.4) and 150 mM NaCl, then treated with lambda phosphatase (New England Biolabs) in 1x NEBuffer for PMP and 1 mM MnCl_2_ and incubated at 30°C for 30 to 60 min with agitation.

For immunoblotting, proteins were resolved by PAGE (see below), transferred to a polyvinylidene difluoride (PVDF) membrane (Immobilon FL; EMD Millipore), blocked with Odyssey Blocking Buffer (LI-COR Biosciences), and incubated with primary antibody at 2 μg/mL or 1:5000 anti-Dma1 serum or 1:2000 anti-Sid4 serum overnight at 4°C. Primary antibodies were detected with secondary antibodies coupled to IRDye680 or IRDye800 (LI-COR Biosciences, Lincoln, NE) and visualized using an Odyssey Infrared Imaging System (LI-COR Biosciences). Resolving gels: 3–8% Tris-acetate PAGE used for Dma1-FLAG and Sid4-DUB blotting; 4–12% NuPAGE used for Dma1-GFP, Sid4 and Dma1 ubiquitination assay blotting, 12% Tris-glycine PAGE used for Cdc2 blotting.

## Acknowledgements

We thank Drs. Ilektra Kouranti and Tyler McKann for contributing to the cDNA cloning of DUBs used in this study, Anna Feoktistova and Liping Ren for outstanding technical support, Maya Igarashi for technical contributions, Dr. Quanwen Jin for GBP plasmids, and Dr. Colin Gordon for DUB plasmids. We are grateful to Rodrigo Guillen, MariaSanta Mangione, Chloe Snider, and Alaina Willet for critical comments on the manuscript. C.M.J. and A.E.J. were supported by the Cellular, Biochemical and Molecular Sciences Training Program (NIH T32GM08554). Z.C.E. and S.N.C. were supported by the Integrated Biological Systems Training in Oncology Program (2T32CA119925). This work was supported by NIH GM112989 to K.L.G.

## Author contributions

Conception and design of work: C.M.J, J.-S. C., A.E.J., S.N.C., and K.L.G; Acquisition of data: C.M.J, J.-S. C., A.E.J., S.N.C., and Z.C.E; Analysis and interpretation of data: all authors; Writing and editing manuscript: all authors.

## Supplementary Figure Legends

**Figure S1.**
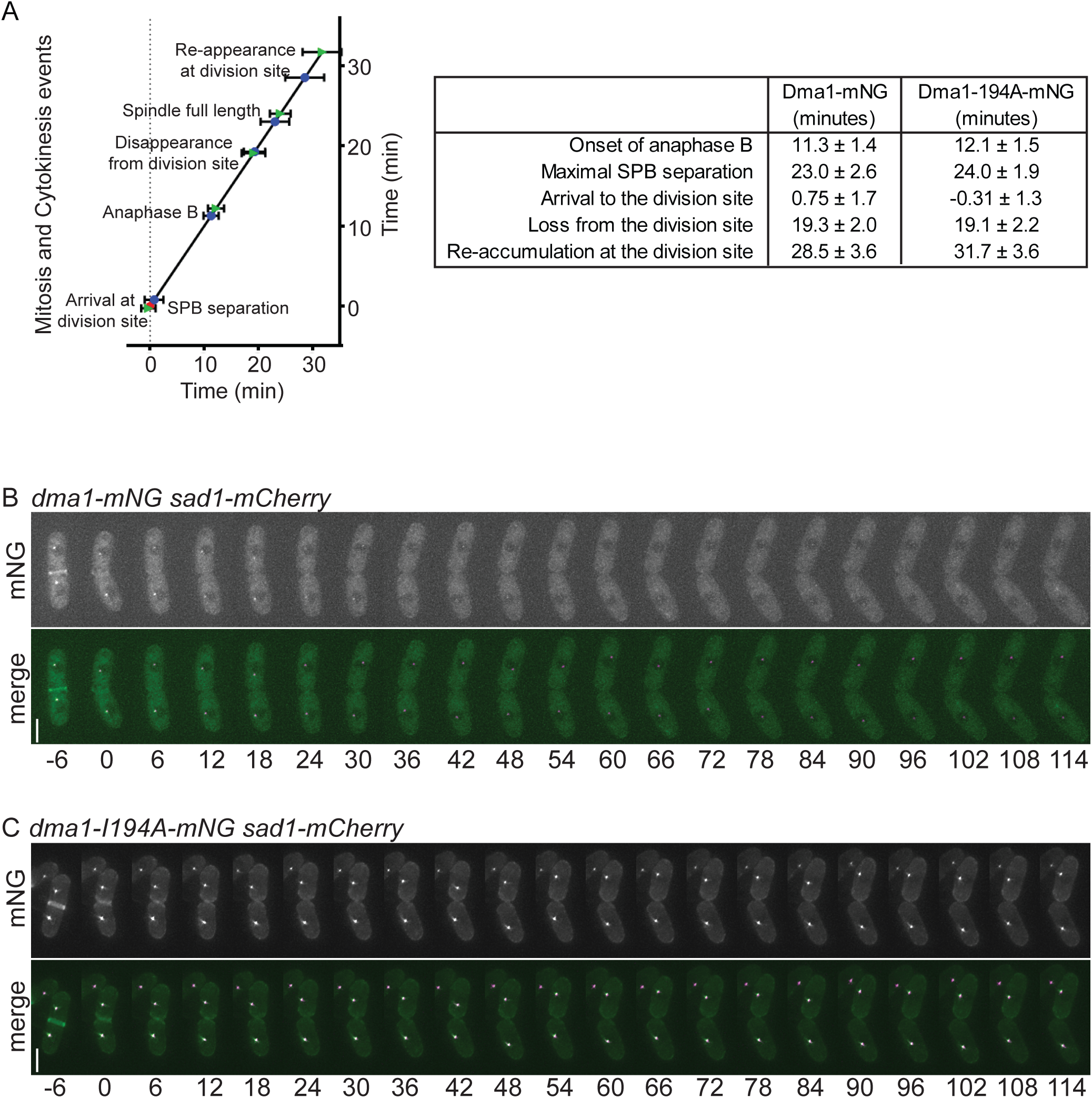
Timing of Dma1 localization relative to mitotic events. (A) (Left, graph) Time line showing the detection of Dma1-mNG (blue circles) and Dma1-I194A-mNG (green triangles) at the division site. Time 0 was defined as SPB separation (red star), and the mean time of detection of each event ± SD is plotted. (Right, table) Average times (minutes) for each strain (top of column) for the onset of anaphase B, maximum SPB separation (end of anaphase B), arrival to the division site, loss from the division site and re-accumulation at the division site. Timing of Dma1-mNG events were determined from 8 cells and Dma1-I194A-mNG events were determined from 13 cells. (B-C) Images from representative movies of dividing *dma1-mNG sad1-mCherry* (B) and *dma1-I194A-mNG sad1-mCherry* (C) cells. Time in minutes denoted below images, 0 indicates initial frame that physical separation of daughter cells began. Scale bars, 5 μm.

**Figure S2.**
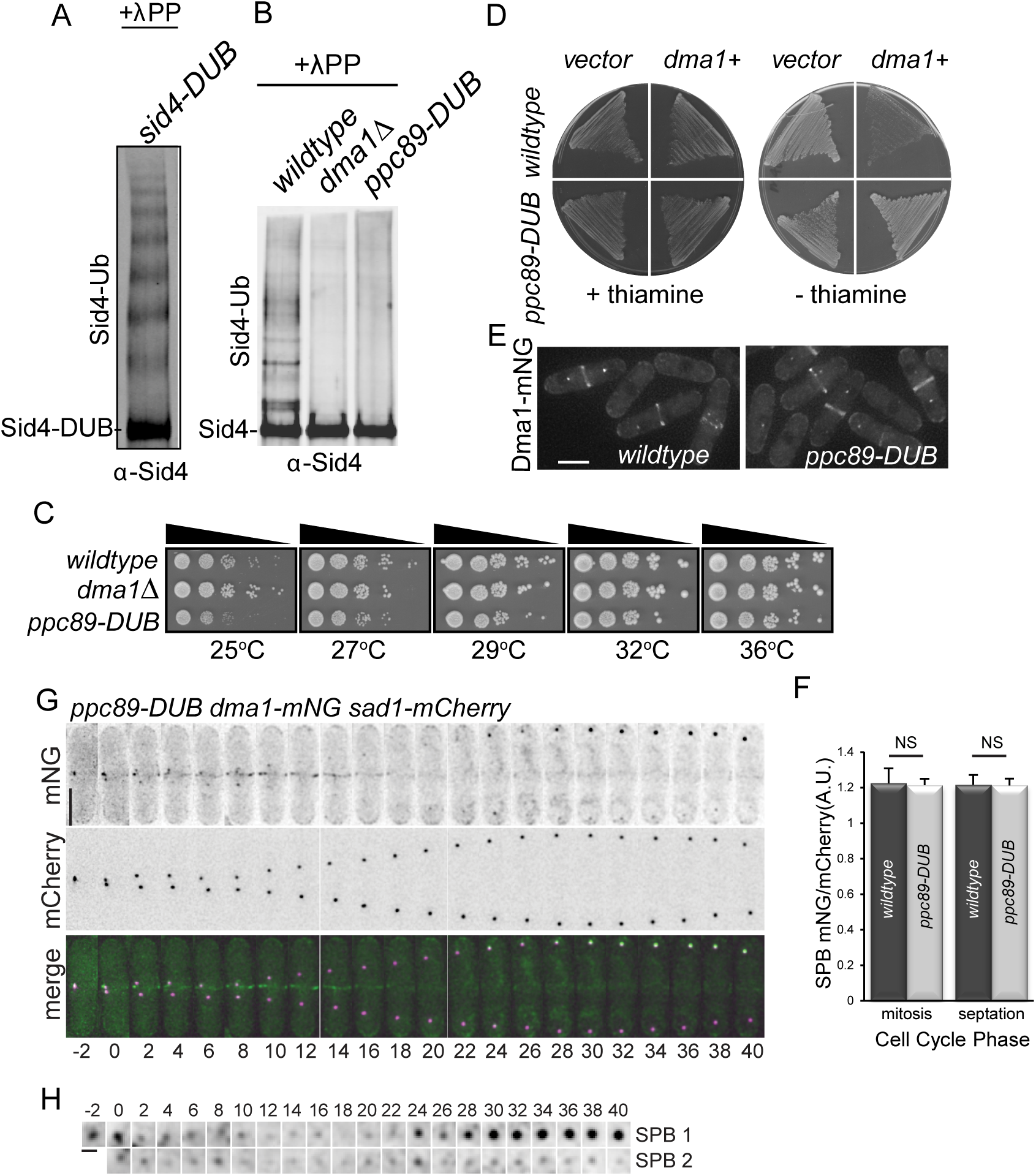
A ppc89-DUB fusion eliminates Sid4 ubiquitination. (A) A Sid4-DUB fusion was immunoprecipitated from the indicated strain, treated with phosphatase, and visualized by immunoblotting. (B) Sid4 was immunoprecipitated from equal amounts of protein lysates from the indicated strains, treated with phosphatase, and visualized by immunoblotting. C) The indicated strains were grown at 29°C, and then the same number of cells were spotted in 10-fold serial dilutions and incubated at the indicated temperatures on YE plates. D) *wildtype* and *ppc89-DUB* strains were transformed with either vector alone (pREP42) or vector containing *dma1^+^* (pREP42dma1^+^). Transformants were incubated on medium containing thiamine (left) or lacking thiamine (right) at 29°C. E) Live cell images of Dma1-mNG in *wildtype* and *ppc89-DUB* cells. Scale bar, 5 μm. F) Quantification of Dma1-mNG at SPBs in the indicated strains, relative to Sad1-mCherry. n ≥ 40 cells for each; error bars represent standard error of the mean, NS = not significant. (G) Images at 2-min intervals from a representative movie of *ppc89-DUB dma1-mNG sad1-mCherry*. Time 0 indicates initial frame of SPB separation. Scale bar, 5 μm. (H) Enlarged SPB region(s) from movie in G. Scale bar, 1 μm.

**Figure S3.**
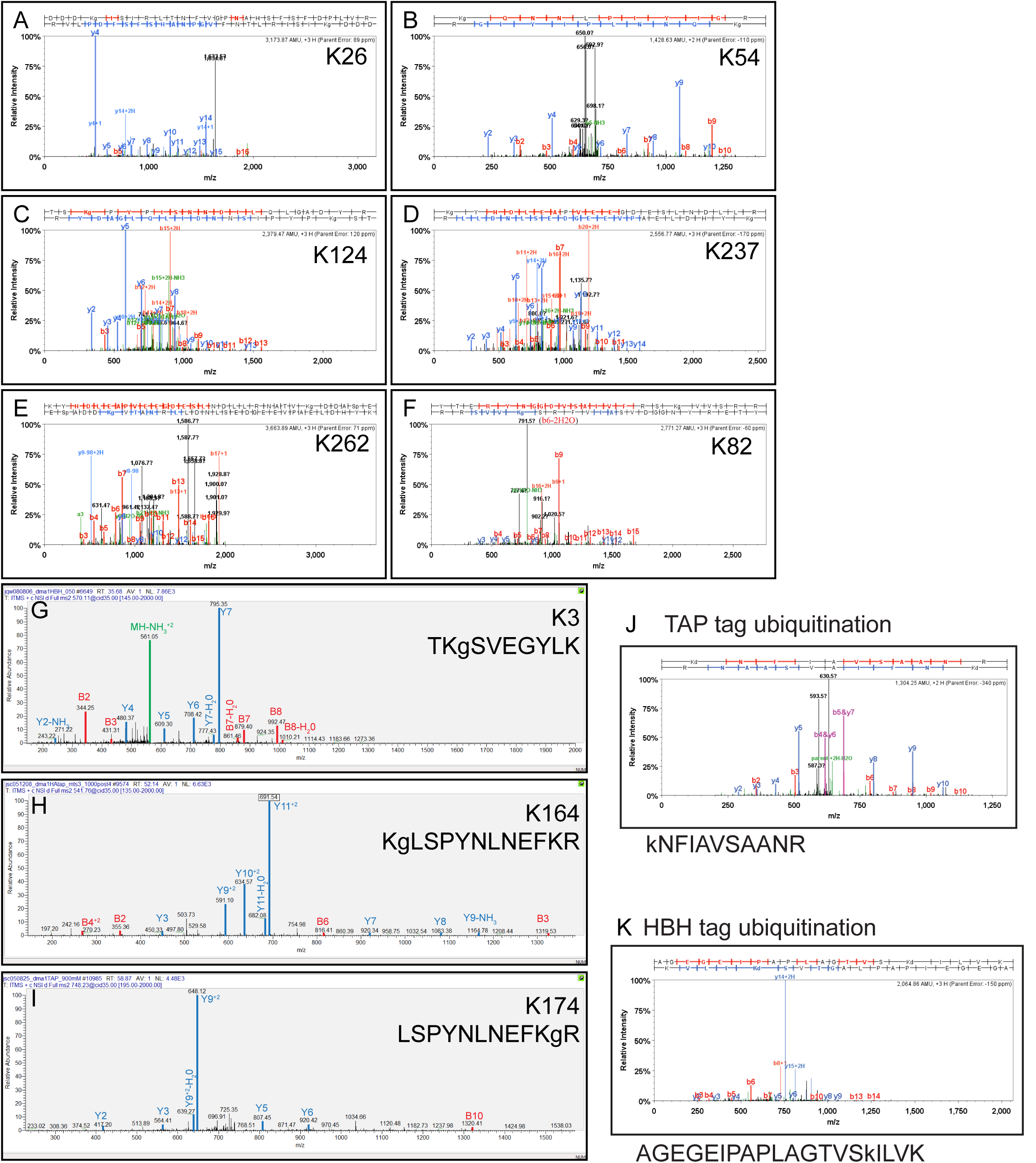
Mass spectra indicative of Dma1 ubiquitination. (A-I) Representative MS^2^ spectra of Dma1 ubiquitination at K26 (A), K54 (B), K124 (C), K237 (D), K262 (E), K82 (F), K3 (G), K164 (H), and K174 (I) and K49 of the TAP tag (J) and K35 of the HBH tag (K). Spectra shown in A-F and J-K were generated and annotated in Scaffold and G-I were identified by Myrimatch, displayed in Xcalibur (v 2.2, Thermo Scientific) and manually annotated. Fragment ions are indicated in each spectrum (y-ions in blue, b-ions red and neutral loss ions in green).

**Figure S4.**
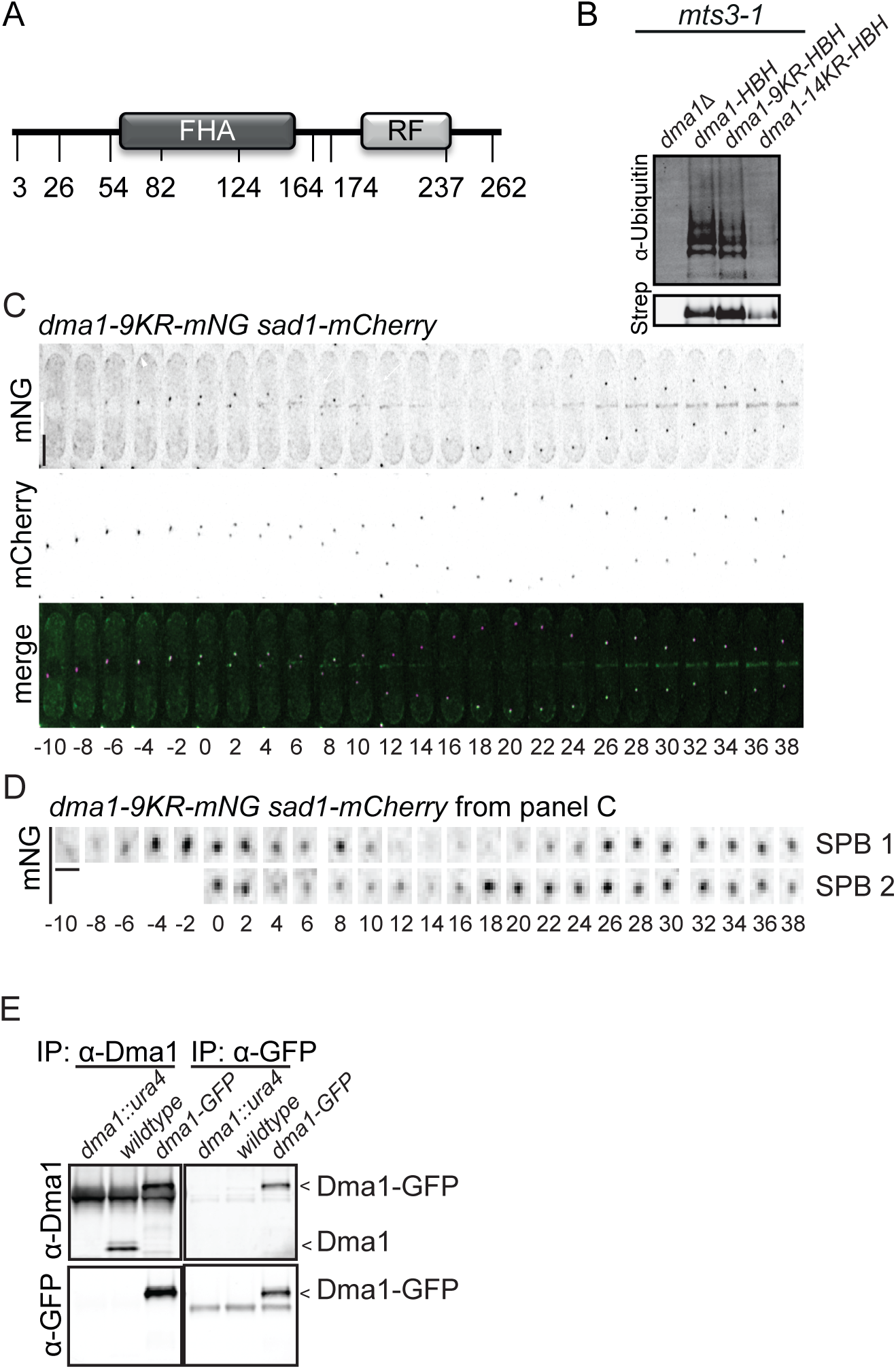
Dma1–9KR-mNG mutant exhibits wildtype localization dynamics. (A) Schematic of Dma1 with ubiquitination sites identified by LC-MS/MS indicated. (B) Dma1-HBH, Dma1–9KR-HBH, Dma1–14KR-HBH, or non-specifically purified proteins were isolated from *mts3–1* cells shifted to 36°C for 3 h. Dma1 ubiquitination was detected by immunoblotting with an anti-ubiquitin antibody (top panel) and unmodified Dma1 was detected with fluorescently-labelled streptavidin (bottom panel). (C) Images from representative movie of *dma1–9KR-mNG sad1-mCherry*. Time in minutes denoted below images; 0 indicates initial frame of SPB separation. Scale bar, 5 μm. (D) Enlarged SPB region(s) from movie in C. Scale bars, 1 μm. (E) Dma1 or Dma1-GFP was immunoprecipitated from the indicated strains with anti-Dma1 serum (left panels) or anti-GFP antibody (right panels). Dma1 and Dma1-GFP were detected by immunoblotting with an anti-Dma1 serum (top panels) or anti-GFP antibody (bottom panels). Dma1 and Dma1-GFP are marked with arrowheads.

**Table S1.**
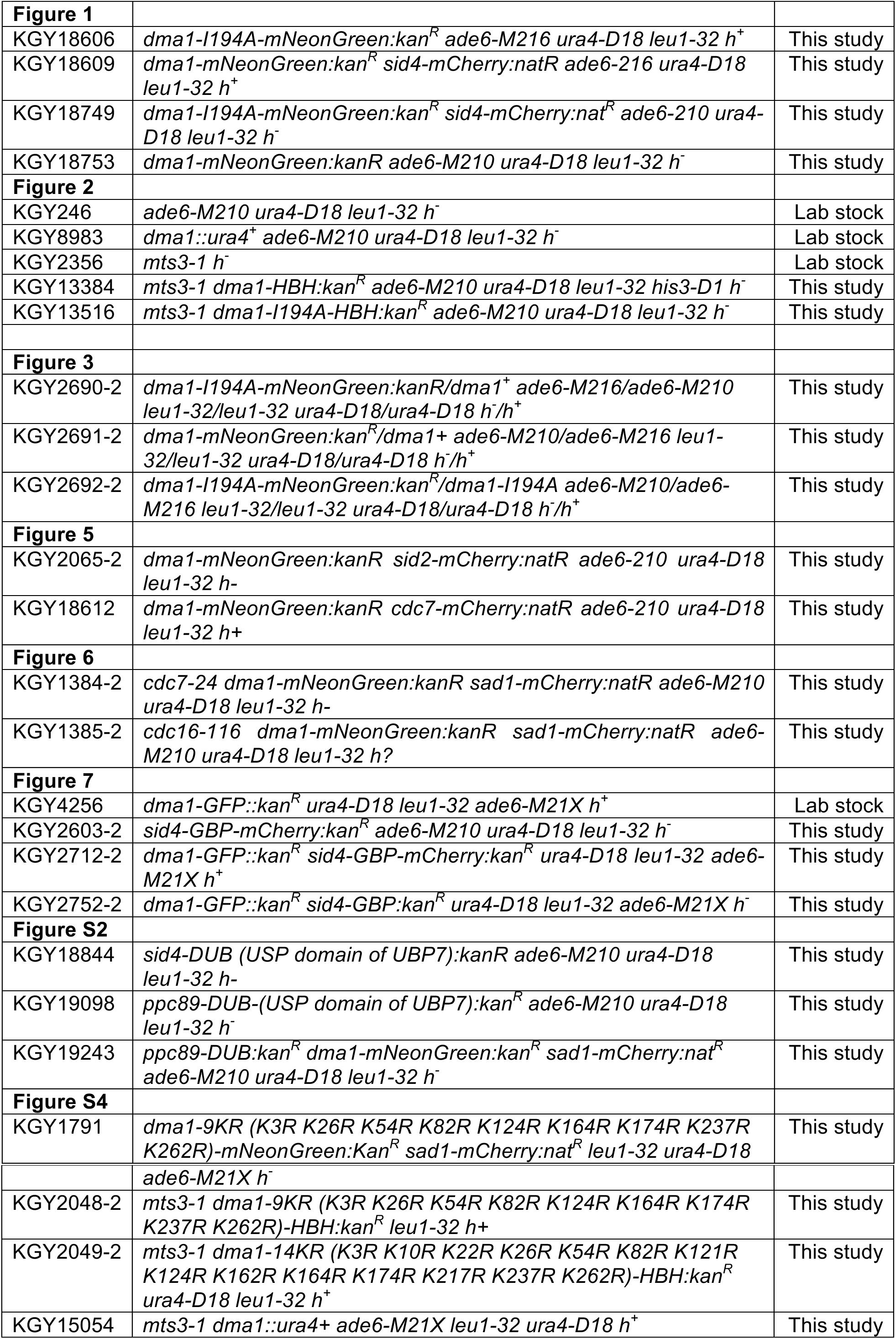
*S. pombe* strains used in this study

